# Sex-specific predictive values of biomarkers for immunotherapy efficacy in lung adenocarcinoma

**DOI:** 10.1101/2020.10.26.356220

**Authors:** Mingming Jia, Tian Chi

## Abstract

It remains a challenge to accurately predict patient responses to tumor immunotherapy, although various biomarkers have been proposed to predict patient responses to anti-PD-1 therapy. Here by integrating genomic, transcriptomic, proteomic and clinical phenotype data from three immunotherapeutic cohorts and a multiple-dimensional dataset of The Cancer Genome Atlas (TCGA) project, we uncovered a profound effect of Sex on the predictive values of conventional biomarkers in lung adenocarcinoma (LUAD): only in females were nonsynonymous mutation burden (TMB), neoantigen burden, smoking signature, KRAS mutations (especially G12C and G12V) or tumor microenvironment robustly correlated with anti-PD-1 efficacy; the correlations in males were either absent or weaker. We propose that Sex be considered in conjunction with conventional biomarkers when predicting immunotherapy efficacy, and conversely, conventional biomarkers be carefully controlled for when attempting to dissect the impact of Sex on immunotherapy efficacy.

## Introduction

PD-1 blockade immunotherapy holds great promises for combating cancers including non-small cell lung cancer (NSCLC) (1–4). Unfortunately, only a minority of NSCLC patients respond to the therapy (1–3, 5, 6). Multiple biomarkers for NSCLC sensitivity to PD-1 blockade have been proposed, including nonsynonymous tumor mutation burden (TMB) (1, 2), neoantigen burden (1, 2), smoking signature (1, 7), KRAS mutation (1, 7), and tumor microenvironment (TME) (7, 8). However, these factors, which seem to function in conjunction (1, 6, 8), are insufficient to accurately predict the immunotherapy efficacy (5, 6, 8).

Sex dimorphism has been documented for human cancers, with males at a higher risk of getting and dying of most types of cancers throughout life (9). Notably, among all cancers, lung cancers display one of the greatest Sex-specific differences (10). For instance, females exposed to cigarette smoke and female never-smokers are each more susceptible to lung cancers than the male counterparts, consistent with the ability of estrogen to stimulate cancer cell proliferation (11). Finally, sex dimorphism in the survival rates has been reported in a mouse model of metastatic lung cancer resulting from TP53 and KRAS mutations (12).

Very little is known about whether Sex affects the outcome of cancer immunotherapy. A recent meta-analysis concludes that immunotherapy is more effective in men in diverse cancers (13), but the argument is controversial (14–17). In addition, better responses to PD-L1 blockade has been observed in female mice with melanoma (18), but whether this is true in humans or for other cancers is unclear.

Here we report that Sex, combining with known biomarkers, indicates the predictive values for anti‑PD-1 therapy efficacy in lung adenocarcinoma (LUAD), one of the most common types of NSCLC (4, 19).

## Materials and Methods

### Three NSCLC cohorts with immunotherapy: *Science*, *JCO* and *Cancer Cell*

We used clinical characteristics and mutation data from three NSCLC datasets, initially described in *Science* (1), *J. Clinical Oncology* (*JCO*) (3) and *Cancer Cell* (2), respectively (Table 1).

**Table 1.** Characteristics and analysis of the 4 datasets used in this study. irRC, immune-related Response Criteria; RECIST, Response Evaluation Criteria in Solid Tumors; WES, Whole-Exome Sequencing; MSK-IMPACT, Memorial Sloan-Kettering Integrated Mutation Profiling of Actionable Cancer Targets; Tv/Ti, Transversion/Transition. ^1^excluded from CR/PR and SD/PD but included in progression-free survival (PFS) assessment; ^2^excluded from DCB/NDB but included in PFS evaluation; ^3^excluded from DCB/NDB analysis. ^4^ mutation information insufficient; ^5^MAF files unavailable; ^6^samples and KRAS mutations too few; ^7^Transcriptom profiles unavailable.

|  | *Science-LUAD* | *Cancer Cell-NSCLC* | *JCO-LUAD* | *TCGA-LUAD* |
| --- | --- | --- | --- | --- |
| Immunotherapy | Yes | Yes | Yes | No |
| No. of patients | 29 | 75 | 186 | 478 |
| Age (years), median (range) | 63(41-80) | 66(42-87) | 66(22-92) | 67(38-88) |
| Gender |  |  |  |  |
| Male | 13(45) | 37(49) | 83(45) | 221(46) |
| Female | 16(55) | 38(51) | 103(55) | 257(54) |
| Smoking status |  |  |  |  |
| Ever | 24(83) | 60(80) | 145(78) | 169(35) |
| Never | 5(17) | 15(20) | 41(22) | 29(6) |
| Unknown | 0(0) | 0(0) | 0(0) | 280(59) |
| Histology |  |  |  |  |
| LUAD(lung adenocarcinoma) | 29(100) |  | 186(100) | 478(100) |
| LUSC(lung squamous cell carcinoma) |  | 16(21) |  |  |
| non-squamous cell carcinoma |  | 59(79) |  |  |
| PD-L1 expression |  |  |  |  |
| 0% | 4(14) | 25(33) | 36(19) |  |
| >=1% | 22(76) | 45(60) | 30(16) |  |
| Unknown | 3(10) | 5(7) | 120(65) | 478(100) |
| Treatment (antibody) | PD-1 | PD-1 + CTLA-4 | PD-(L)1 |  |
| Method for efficacy evaluation | irRC | RECIST v1.1 | RECIST v1.1 |  |
| Best overall response |  |  |  |  |
| Complete/Partial Response (CR/PR) |  | 24(32) |  |  |
| Stable/Progressive Disease (SD/PD) |  | 44(59) |  |  |
| Not evaluable (<6 months follow-up) |  | 7(9)<sup>1</sup> |  |  |
| Clinical benefit |  |  |  |  |
| Durable clinical benefit (DCB) | 12(41) |  | 55(30) |  |
| No durable benefit (NDB) | 15(52) |  | 124(66) |  |
| Not evaluable (<6 months follow-up) | 2(7)<sup>2</sup> |  | 7(4)<sup>3</sup> |  |
| Mutation detection method | WES | WES | MSK-IMPACT | WES |
| Transcriptom profiling |  |  |  | RNA-seq |
| Biomarker predictive value (gender mixed) |  |  |  |  |
| Tumor mutational burden (TMB) | Yes | Yes | Yes | Yes (ref. 1,7,22) |
| Neoantigen burden | Yes | Yes | Not done | Yes (ref. 25,32) |
| Smoking signature (Tv/Ti) | Yes | No | No | Yes (ref. 1, 7) |
| Smoking signature (SignatureAnalyzer) | Not done | Not done | Not done | Yes(ref. 22,28) |
| KRAS mutation | Yes | No | No | Yes (ref. 4,7) |
| Tumor microenvironment | Not done | Not done | Not done | Yes (ref.7,25,30) |
| **Biomarker predictive value (robust only in female, reported in this study)** |  |  |  |  |
| Tumor mutational burden (TMB) | Yes (Fig. 1) | Yes (Fig. 1) | Yes (Fig. S2) | Fig. S5 |
| Neoantigen burden | Yes (Fig.2) | Yes (Fig.2) | Unknown<sup>4</sup> | Fig. S5 |
| Smoking signature (Tv/Ti) | Yes (Fig. S4) | Yes (Fig. S4) | Yes (Fig. S4) | Yes (Fig. S4) |
| Smoking signature (SignatureAnalyzer) | Yes (Fig.3) | Not done<sup>5</sup> | Not done<sup>5</sup> | Yes (Fig.3) |
| KRAS mutation | Unknown<sup>6</sup> | Yes (Fig.4) | Yes (Fig. 4) | Yes (Fig.4) |
| Tumor microenvironment | Not done<sup>7</sup> | Not done<sup>7</sup> | Not done<sup>7</sup> | Yes (Fig.5-6; S7) |

The *Science* dataset contains 34 NSCLC (29 LUAD) patients on a pembrolizumab (anti-PD-1) therapy initiated in 2012-2013 (protocol NCT01295827). The therapeutic effects were classified into “durable clinical benefit” (DCB, with partial response /stable diseases lasting > 6 months) and “no durable benefit” (NDB, the benefits persisting < 6 months), based on investigator-assessed immune-related Response Criteria (irRC). Mutations were detected using whole exome sequencing (WES) on the Illumina platform.

The *JCO* dataset involves 240 NSCLC (186 LUAD) patients on anti-PD-1 and/or anti-PD-L1 monotherapies. The therapeutic effects were classified into DCB/NDB based on Response Evaluation Criteria in Solid Tumors (RECIST) guideline, version 1.1. Mutations were mapped for 468 genes using MSK-IMPACT targeted-sequencing.

The *Cancer Cell* dataset involves 75 NSCLC patients on nivolumab plus ipilimumab (to block PD-1 and CTLA-4) initiated in 2013-2015 (protocol NCT01454102). The therapeutic effects were classified into Complete Response/Partial Response (CR/PR) and Stable Disease/Progressive Disease (SD/PD) using RECIST version 1.1. Mutations were detected using whole exome sequencing (WES) on the Illumina platform. Of note, the *Cancer Cell* cohort comprises non-squamous cell carcinoma (79%, 59/75) and squamous cell carcinoma (21%, 16/75), but adenocarcinoma (LUAD), one of the commonest NSCLC types, is not explicitly marked. We therefore included all 75 NSCLC samples in our analysis. As such, the *Cancer Cell-NSCLC* cohort has a caveat and is only peripheral to our study. In addition, we note that whereas the patients in the *Science* and *JCO* cohorts were treated with PD-(L)1 blockade, PD-1 plus CTLA-4 blockade was used instead in the *Cancer Cell* cohort (Table 1), which might help explain its unique therapeutic responses, given the differences of molecular mechanism in PD-1 vs. CTLA-4 blockade (20).

### Clinical cohorts without immunotherapy: *TCGA-LUAD* datasets

The *TCGA-LUAD* dataset was collected from https://xenabrowser.net/datapages/, which include profiles on somatic mutations (n=543), mRNA expression (n=576), and protein expression (n=365). Mutations were detected using whole exome sequencing (WES) on the Illumina platform, with the calls calculated at Broad Institute Genome Sequencing Center, and only the calls with variant allele frequency (VAF) above 4.0% kept. RNA-seq was done using the Illumina HiSeq 2000 RNA Sequencing platform. Briefly, gene-level transcription was estimated as log2(x+1) transformed RSEM normalized counts, with the units of mRNA expression values being pan-cancer normalized log2 (norm_count+1). We removed adjacent normal tissues and only focused on primary LUAD samples (n=478, three samples whose WES-matched expression profiles were missing). Finally, protein expression was profiled at the MD Anderson Cancer Center TCGA proteome characterization center RPPA core. The units of protein expression values were normalized RPPA (reverse phase protein array) values. The detailed RPPA method has been previously described (19).

### Receiver Operator Characteristic (ROC) curves and Youden’s index

We used ROC curves to plot the true-positive rate (sensitivity) against the false-positive rate (1 − specificity) at various threshold settings. ROC plots showed the tradeoff between sensitivity (ability to detect DCB or CR/PR patients) and specificity (ability to detect NDB or SD/PD patients) when creating a diagnostic test. The area under a ROC curve (AUC) quantified the overall ability of the test to discriminate between those individuals with DCB or CR/PR and those with NDB or SD/PD. A truly useless diagnostic test has an AUC of 0.5, and a perfect test has an AUC of 1.0. Youden’s index (equaling sensitivity + specificity – 1) is a summary measure of the performance of a ROC curve, and the maximum value of the index was used as a criterion for choosing the optimum cut-off point for TMB and neoantigen burden (1, 21). These cut-off points were then used to divide the patients into high vs. low TMB/neoantigen burden categories. Note that ROC tests performed much better in females, with greater AUC and Youden’s index. Thus, the index–associated cutoffs for TMB and neoantigen burden were more reliable for predicting therapy efficacy in females.

### Gene Set Enrichment Analysis (GSEA)

We performed gene differential expression analysis and heat maps execution using Multiple Experiment Viewer (MeV; http://mev.tm4.org/#/welcome). Sample clustering was done using MeV hierarchical unsupervised clustering function with Pearson’s correlation distance and average linkage (differential expression genes with FDR-adjusted *p* value ≤0.005). For GSEA, MsigDB (Molecular Signatures Database) (22) was used to enrich the significant pathway, with q value less than 0.05 considered statistically significant.

### Immune cellular infiltration estimates

We captured CIBERSORT algorithm (23) to calculate the infiltration of immune cells in the TME. Using influential deconvolution method, CIBERSORT could estimate the abundance of immune cells in each tumor sample based on gene expression data, and the CIBERSORT analysis results of *TCGA-LUAD* samples could be acquired from the Pan-Cancer Atlas (24). These cell types include total macrophages, various macrophage subsets (M0, M1 and M2), total lymphocytes and various subsets thereof (B cell naïve, B cell memory, CD4 T naïve, CD4 T cell memory resting, CD4 T cell memory activated, T cell follicular helper, Tregs, T cell gamma-delta, CD8 T cells, NK cell resting, NK cell activated, and Plasma cells) (24).

### Mutational landscape analysis

To calculate the mutational landscape, we first extracted *TCGA-LUAD* level 3 Mutation Annotation Format (MAF) files from http://gdac.broadinstitute.org/. We then used “maftools” R packages (25) to calculate total somatic nonsynonymous mutation counts and the percentages of KRAS nonsynonymous mutations. KRAS mutation spectrum of other three immunotherapeutic cohorts were extracted from the corresponding supplemental materials in the original publications. Finally, the percentages of each of six possible single nucleotide changes within the codons (C>A:G>T, C>G:G>C, C>T:G>A, T>A:A>T, T>C:A>G, T>G:A>C) were calculated to generate a 96-feature vector, which was used to represent the mutation spectrum in that sample.

### *In silico* neoantigen prediction

Neoantigen information for *Science-LUAD* and *Cancer Cell-NSCLC* cohorts were downloaded from the Supplemental Materials in the original publications (1, 2), while that of the *TCGA-LUAD* cohort acquired from the Pan-Cancer Atlas (24). We excluded the *JCO-LUAD* cohort because its mutation profiles were based on MSK-IMPACT targeted-sequencing and available for only 468 genes (26), which is insufficient for neoantigen calculation. The neoantigen prediction pipelines were divergent between *Science-LUAD* and *Cancer Cell-NSCLC* (1, 2): in the former, mutated DNA was translated using NAseek and the MHC-I binding affinities of the resulting peptides predicted using NetMHCv3.4; for the latter, nonsynonymous point mutations were translated using Topiary and affinities predicted with NetMHCcons (v. 1.1).

### Smoking signature analysis

For mutation signature analysis, we captured R package “SignatureAnalyzer” (27, 28) that could use non-negative matrix factorization algorithm (NMF) to decipher and calculate mutation signatures based on cancer somatic mutations in tri-nucleotide sequence contexts (https://software.broadinstitute.org/cancer/cga/msp). “SignatureAnalyzer” could then automatically generate the mutation signatures (W) and signature activities across samples. MAF files are necessary for this analysis and available in the *Science-LUAD* and *TCGA-LUAD* datasets, which were therefore analyzed.

### Statistical Analyses

Statistical analyses were performed using R software (version 3.5.2) and GraphPad Prism (version 7.0.0). For values that approximate normal distribution, Student’s t test or one-way ANOVA was used to determine the differences; otherwise, Mann-Whitney U test or Kruskal-Wallis test was used. The log-rank test was used to compare Kaplan-Meier survival curves of progression free survival (PFS) and the log-rank method was used to determine hazard ratios. The proportion of KRAS mutation was compared using Fisher’s exact test. Pairwise correlations among TMB, neoantigen burden, Tv/Ti, and PFS were calculated using Pearson correlation formula. Logistic and Cox regression were conducted to assess the impact of TMB on clinical response and PFS, respectively, while adjusting for other covariates described. All reported P values were two-tailed, and for all analyses, P≤0.05 was considered statistically significant.

## Results

### Study design

To explore the potential effect of Sex on the predictive values of conventional biomarkers of immunotherapy efficacy, we focused on lung cancers, where multiple clinical trials have been conducted (4). We searched the literature for datasets that provide information on cancer genomes and corresponding responses to immunotherapy. Three non-small cell lung cancer (NSCLC) cohorts (published in *Science* (1), *JCO* (3) and *Cancer Cell* (2)) were found. Multiple potential biomarkers have been examined in these cohorts, with some (TMB and neoantigen) consistently demonstrating predictive values while others (smoking signature and KRAS mutations) informative only in a subset of the cohorts; Table 1). Our strategy was to re-analyze the datasets essentially as described in the previous studies except that we stratified the patients according to Sex.

The sample sizes in the three immunotherapy datasets are small. Fortunately, a multi-dimensional tumor profiling dataset with a much larger sample size (*TCGA-LUAD*) is available that does not involve immunotherapy but can nevertheless allow the inference of biomarkers for immunotherapy responses (7, 24, 29–31). We therefore integrated *TCGA-LUAD* with the three immunotherapeutic datasets to examine how Sex affects the performance of each of the 5 commonly used immunotherapy biomarkers (1, 7, 8) (TMB, neoantigen burden, smoking signature, KRAS mutations and tumor microenvironment) in LUAD. Note that among the 4 datasets, only *TCGA-LUAD* contains (sufficient) information on all the biomarkers, while the other three fit for the analysis of only subsets of the biomarkers (Table 1). Using methods essentially identical to those described in previous studies, we validated previous conclusion that the 5 biomarkers could predict therapeutic efficacy (not shown), but report now that these markers were robust only in females (cyanine-shaded in Table 1), as detailed below.

### TMB: analysis of three immunotherapy cohorts

TMB is perhaps the most important biomarker for immunotherapy (32). In *TCGA-LUAD*, *Science-LUAD* and *Cancer Cell-NSCLC*, TMB has been identified using whole exome sequencing (WES), which produced similar numbers of counts across the three datasets (Fig. S1, top). The TMB distribution of these cohorts was skewed to the right, which we resolved using log transformation (Fig. S1, bottom). In contrast to the two datasets above, *JCO-LUAD* contains sequencing data for only 468 genes (3) and therefore features a much lower mutation load per tumor, making it suboptimal for the analysis of TMB and neoantigen burden (Table 1). We therefore presented the analysis of *JCO-LUAD* as supplemental data (Fig. S2)

TMB has been linked to anti-PD-1 efficacy in LUAD (1–3). However, this proved true only for females, as revealed by three types of analyses (Materials and Methods) performed on *Science-LUAD* and *Cancer Cell-NSCLC* (Fig. 1)

**Fig. 1.**
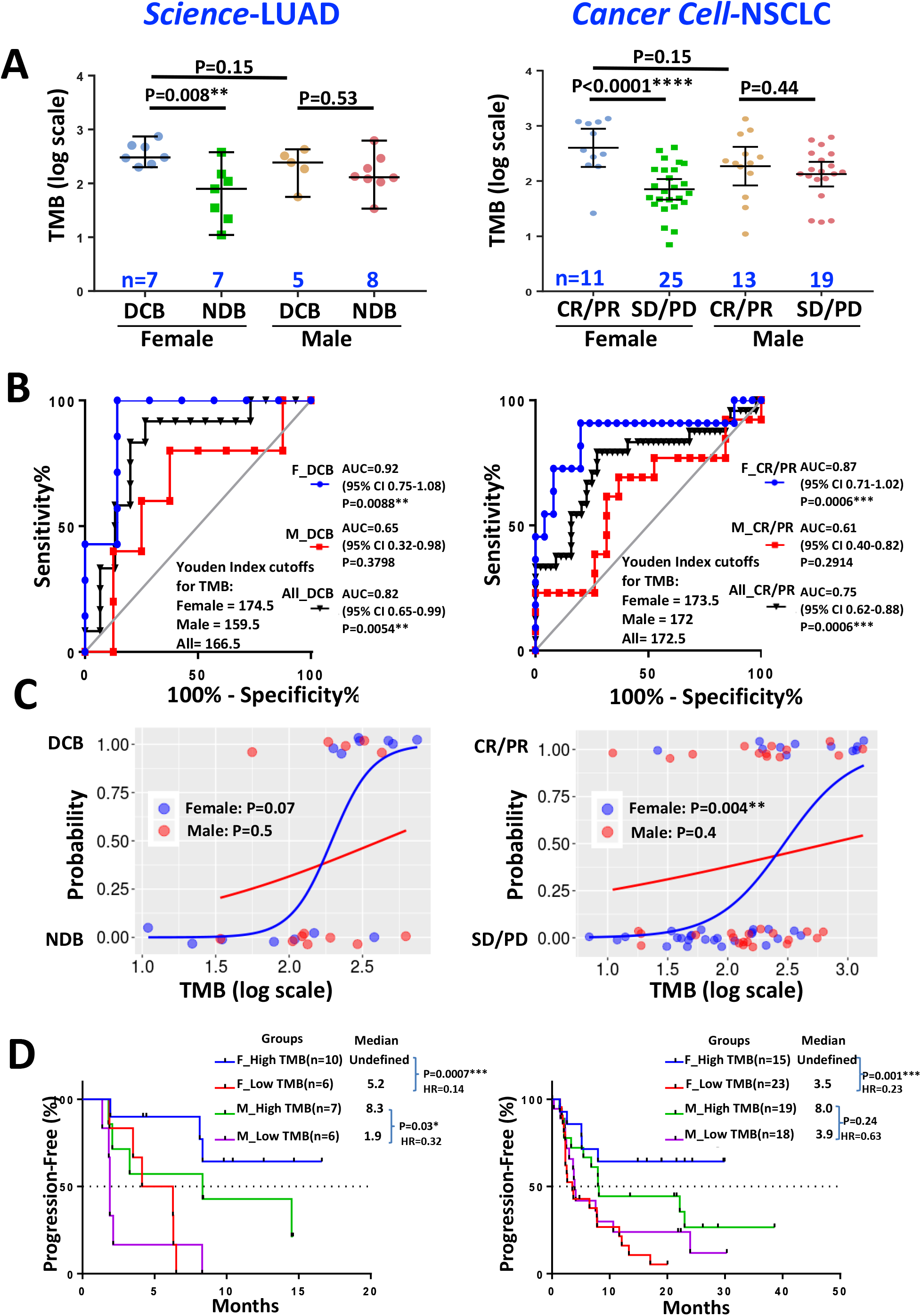
TMB predicted anti-PD-1 efficacy only in females. **(A-C)** Correlation of patient responses to TMB as revealed by scatter plots (A), ROC (B) and logistic regression (C). DCB, durable clinical benefit; NDB, no durable benefit; CR/PR, complete response/partial response; SD/PD, stable disease/progressive disease. AUC, Area under the ROC curve; CI, confidence interval; Probability, probability of getting clinical benefits from immunotherapy. **(D)** Progression-free survival in patients stratified into 4 groups according to Sex and TMB, where High TMB and low TMB were defined as the TMB with values >= and < the Youden Index cutoffs, respectively. HR, hazard ratios.

First, using scatter plots, we found that in females but not males, TMB was higher in patients with better outcomes (Fig. 1A).

Second, ROC analysis was used to measure the true-positive against the false-positive rates at various thresholds of TMB, with the area under the ROC curve (AUC) taken as the quality metric of prediction (1–3, 21). AUC was much higher in the female responders than the male counterparts (0.92 vs. 0.65 and 0.87 vs. 0.61 for the two datasets, respectively; Fig. 1B), demonstrating a strong TMB-prognosis association only in females.

Finally, we used logistic regression to assess the relationship between the TMB and the probabilities of binary therapeutic outcomes (2, 21). S-shaped curves proved a better fit for the data points only in females, supporting the female-specific TMB-prognosis association, although the association in *Science-LUAD* was not statistically significant (p=0.07), presumably reflecting its smaller sample size (n=14; Fig. 1C, left).

The same trend was observed in *JCO-LUAD*, although the Sex effect appeared less obvious (Fig. S2).

We next used the Youden index-based cutoff values (1, 21) for TMB to explore, in *Science-LUAD* and *Cancer Cell-NSCLC*, the relationship between TMB and Progression Free Survival (PFS). The cutoff values were very close in females in the two datasets (174.5 and 173.5, respectively; Fig. 1B). We stratified the females and males each into 2 groups based on their respective cutoff values, and found that in both datasets, high TMB predicted better PFS only in females (Fig. 1D).

Taken together, these data suggest that for LUAD, TMB could reliably predict responses to PD-1 blockade only in females.

### Neoantigen burden: analysis of *Science-LUAD* and *Cancer Cell-NSCLC*

Neoantigens are formed via somatic mutations and can affect therapeutic effect of PD-1 blockade (1, 33). In both *Science-LUAD* and *Cancer Cell-NSCLC*, neoantigen burden predicted therapeutic efficacy in females but not males (Fig. 2) just as TMB did, and not surprisingly, neoantigen burden was significantly correlated with TMB (Fig. S3). Of note, Youden index-based cutoff values of neoantigen burden differed between the two datasets, presumably reflecting the differences in the neoantigen prediction algorithms used in the two studies (See Materials and Methods; Fig. 2B) (1, 2).

**Fig. 2.**
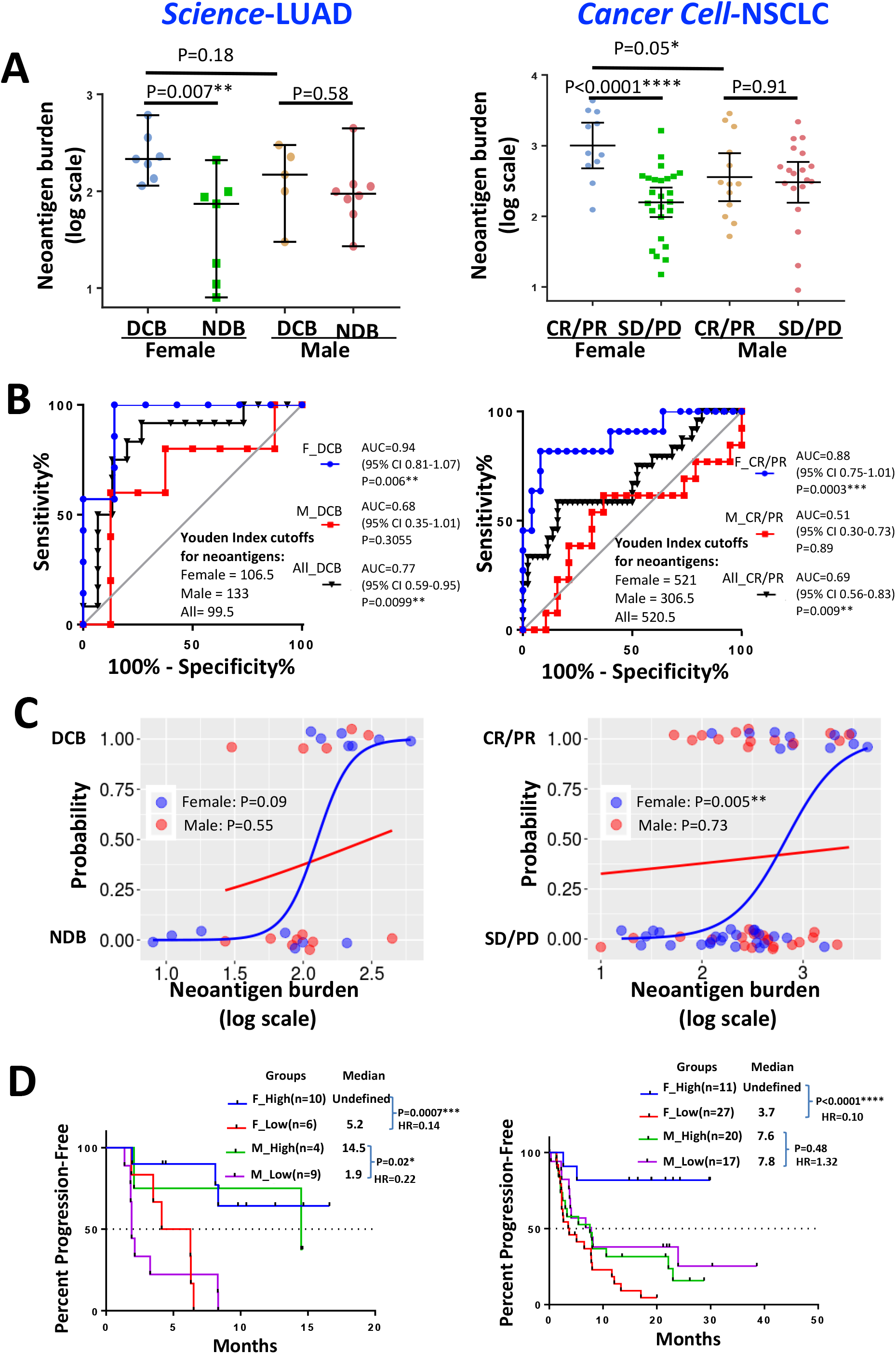
Neoantigen burden predicted anti-PD-1 efficacy only in females. The samples and the methods of their analysis were identical to that in Fig. 1 except that neoantigen burden instead of TMB was examined.

Thus, just as TMB, neoantigen burden predicted therapeutic outcomes in LUAD only in females.

### Smoking signature: analysis of all 4 datasets

A mutational signature is a combinations of mutations characteristic of a specific mutagenesis process (27). Two methods for assessing smoking-related signatures in lung cancer have been reported. The first one, based on simple enumeration of overall transversion (Tv) and transition (Ti) frequencies, is an increase in the Tv/Ti ratio (1, 7, 19). The second smoking-related mutational signature, termed “Signature 4”, has been discovered using SignatureAnalyzer (see Material and Methods) (27, 28). “Signature 4” is characterized by a combination of C>A transversion and, to a less extent, C>T transition, and is probably an imprint of the bulky DNA adducts generated by tobacco smoke and their subsequent removal by non-excision repair (27). Most individual cancer genomes are known to exhibit more than one mutational signatures.

We first analyzed Tv/Ti ratio. In the three immunotherapy cohorts, we found Tv/Ti increased in patients with better therapeutic responses, but only in females (Fig. S4A). In *TCGA-LUAD*, we found similar evidence through examination of the association of Tv/Ti with TMB, and by inference, with patient responses. Briefly, to explore Tv/Ti-TMB association in *TCGA-LUAD*, we stratified the *TCGA-LUAD* samples based on Youden Index cut points for TMB; the cut points were acquired from *Science-LUAD* because it is more homogeneous to *TCGA-LUAD* (Fig. S5A). TMB-Tv/Ti correlation was detected in both sexes, but more robust in females (Fig. S4B). Of note, TMB also correlated with neoantigen burdens, as expected (Fig. S5B-C).

We next used SignatureAnalyzer to examine the smoking signature in *Science-LUAD* and *TCGA-LUAD*, where the MAF files are available (Table 1). We detected two signatures in females with DCB in *Science-LUAD*: W1 (featuring a combination of C>A and C>T mutations) and W2 (featuring C>T mutation; Fig 3A, top), which were related to Signature 4 (a smoking-associated signature) and mismatch repair (MMR) deficiency, respectively (27). W1 was the major signature in these patients, with 5 out of 7 cancer samples displaying only W1 while the remaining 2 a mixture of W1 and W2 (Fig. 3B, top). In contrast, Signature 4-like pattern was depleted in NDB females (whose major signature was related to MMR deficiency; Fig. 3A, bottom and Fig. 3B, bottom), but predominant in males regardless of their therapeutic outcomes (Fig. S6). In *TCGA-LUAD*, Signature 4-like pattern was also preferentially enriched in females with high TMB (compared F_High with F_Low or M_High; Fig. 3C, left) even though these females smoked less than the male counterparts (Fig. 3C, right).

**Fig.3.**
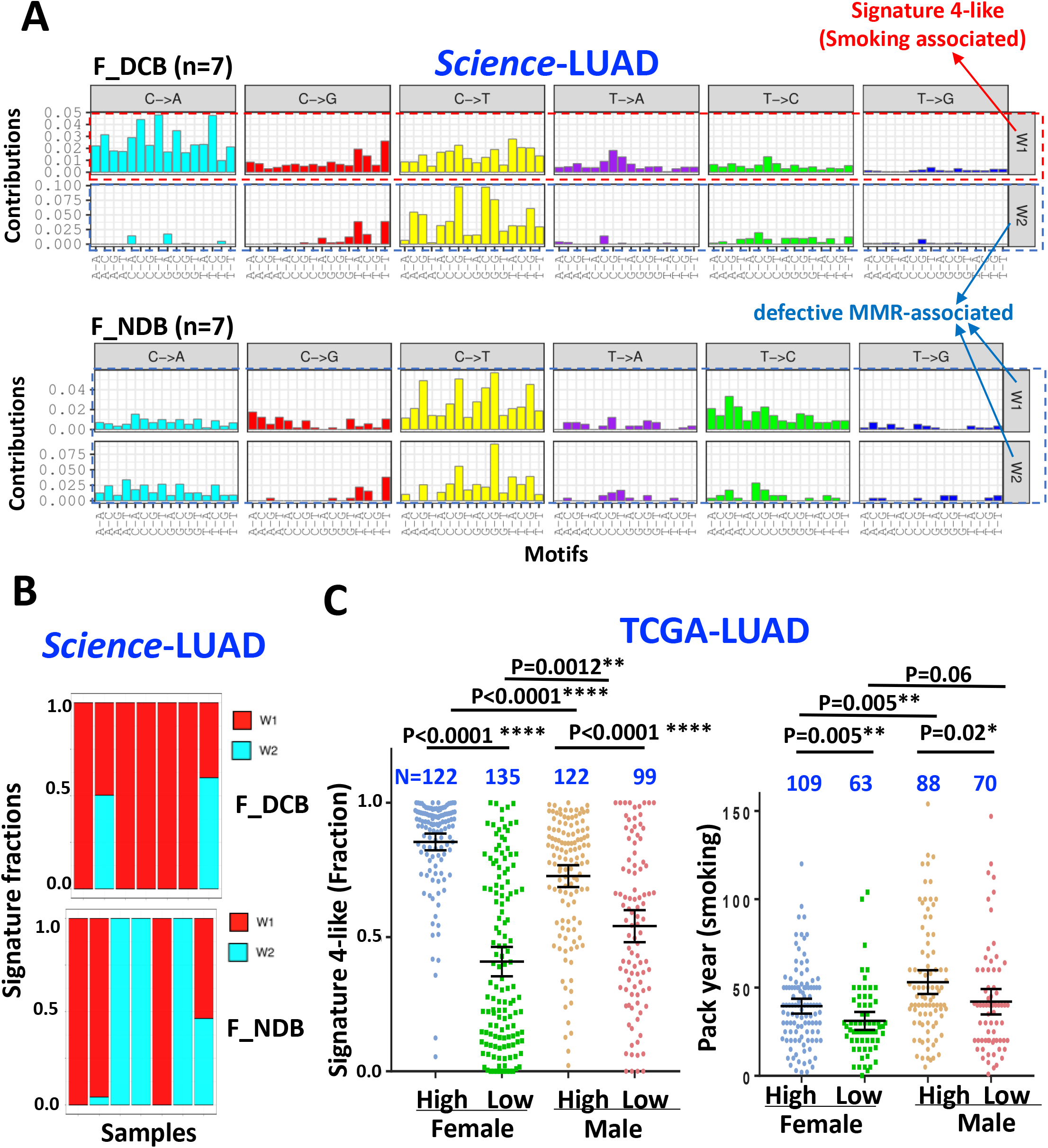
Mutational signature detected using SignatureAanalyzer. **(A)** Mutation spectrum in *Science-LUAD* displayed as the mutated pyrimidines in the context of flanking bases. The probability bars for the six types of substitutions are displayed in different colors. **(B)** Mutation signature composition in individual samples in *Science-LUAD* **(C)** Signature 4-like mutation (left) and smoking doses (right) in various patient groups in *TCGA-LUAD*; patients were classified according to Sex and TMB.

Taken together, these data reinforce the notion that females are more susceptible to tobacco-induced mutations, and suggest that smoking signature (Tv/Ti and Signature 4-like) could robustly predict therapeutic efficacy for LUAD only in female.

### KRAS mutations: analysis of *Cancer Cell-NSCLC*, *JCO-LUAD* and *TCGA-LUAD*

KRAS is often mutated in LUAD, with the G12C/G12V subtypes (resulting from G>T and G>C transversions) being smoking-related (4, 19, 34). The following lines of evidence suggest that KRAS mutations could predict immunotherapy outcomes, but only in females.

First, only in females were overall KRAS mutations and G12C/G12V correlated with therapeutic efficacy (in *Cancer Cell-NSCLC* and *JCO-LUAD*; Fig. 4A, left and middle) and with TMB (in *TCGA-LUAD*; Fig. 4A, right).

**Fig.4.**
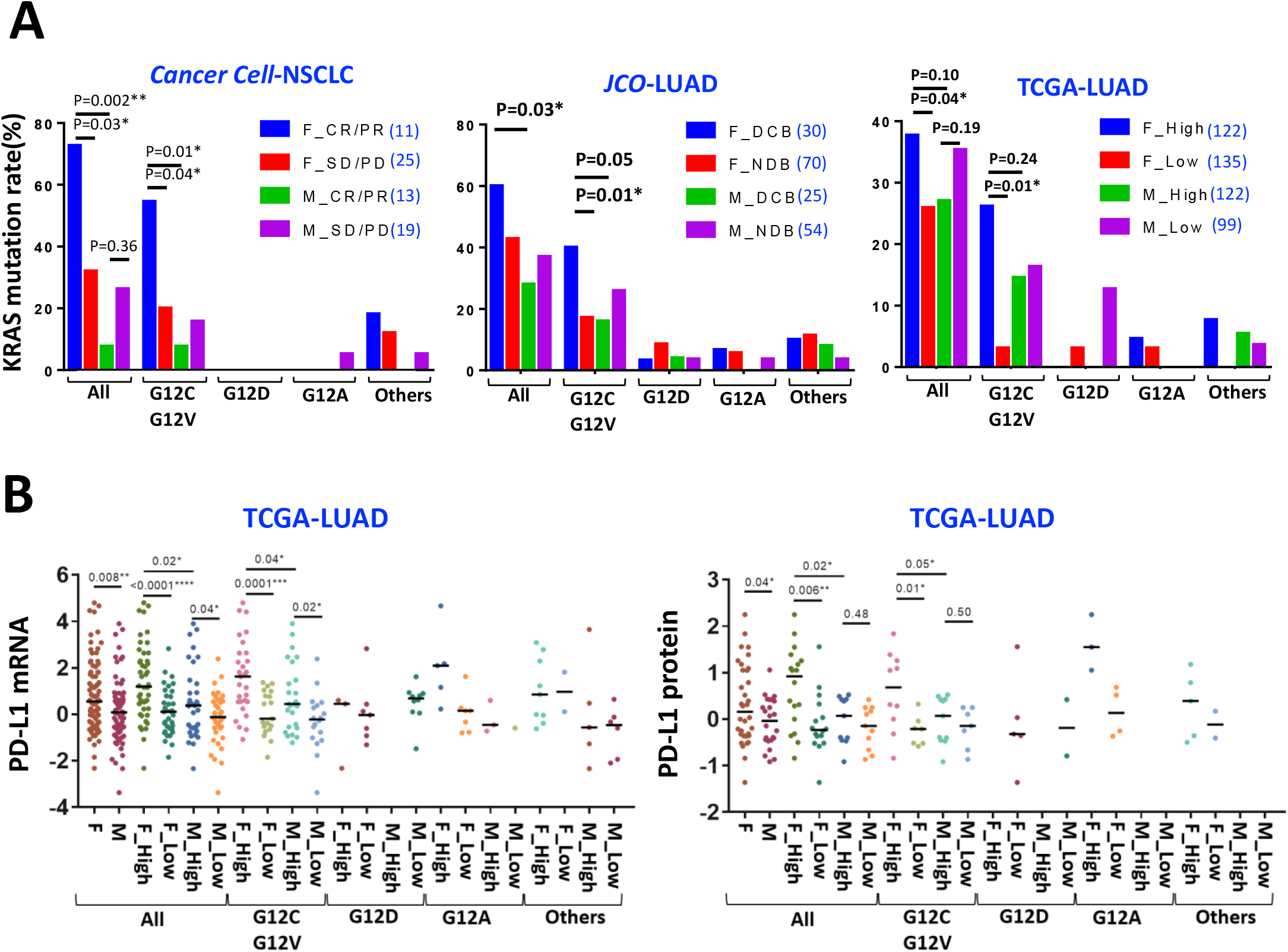
KRAS mutations and their correlations with PD-L1 expression in various patient groups. **(A)** Frequencies of total and various subtypes of KRAS mutations in males and females stratified based on therapeutic responses (left and middle) or TMB (right). *Science*-LUAD is excluded from the analysis for insufficient KRAS mutation counts. Sample numbers are listed next to the sample names. **(B)** Correlation of KRAS mutations with PD-L1 mRNA (left) and protein (right) expression in males and females stratified on TMB.

Second, as revealed in *TCGA-LUAD*, only in females were overall KRAS mutations and G12C/G12V correlated with expression of PD-L1 (Fig. 4B) and immune-related genes (Fig. S7), and by inference, with patient outcomes. Of note, the PD-L1 upregulation might be mediated by KRAS-mediated ERK signaling (35).

### Association of TME with TMB, and by inference, with therapeutic efficacy: analysis of *TCGA-LUAD*

We analyzed gene expression signature and tumor infiltrating cells in TME. We first compared PD-L1 expression among the four patient groups (namely males and females with high or low TMB) and found highest PD-L1 expression in females with high TMB (Fig. 5A). To confirm and extend this observation, we used unsupervised hierarchical clustering to profile differentially expressed genes. Only in females with high TMB could we detect a signature predictive of favorable therapeutic outcomes (7, 29, 36), which is characterized by co-expression of PD-L1 with the immune response genes related to T effector cells and IFN-γ (such as GZMB, CD8A, IFN-γ, CXCL10, CXCL9, GBP1, and STAT1; Fig. 5B).

**Fig.5.**
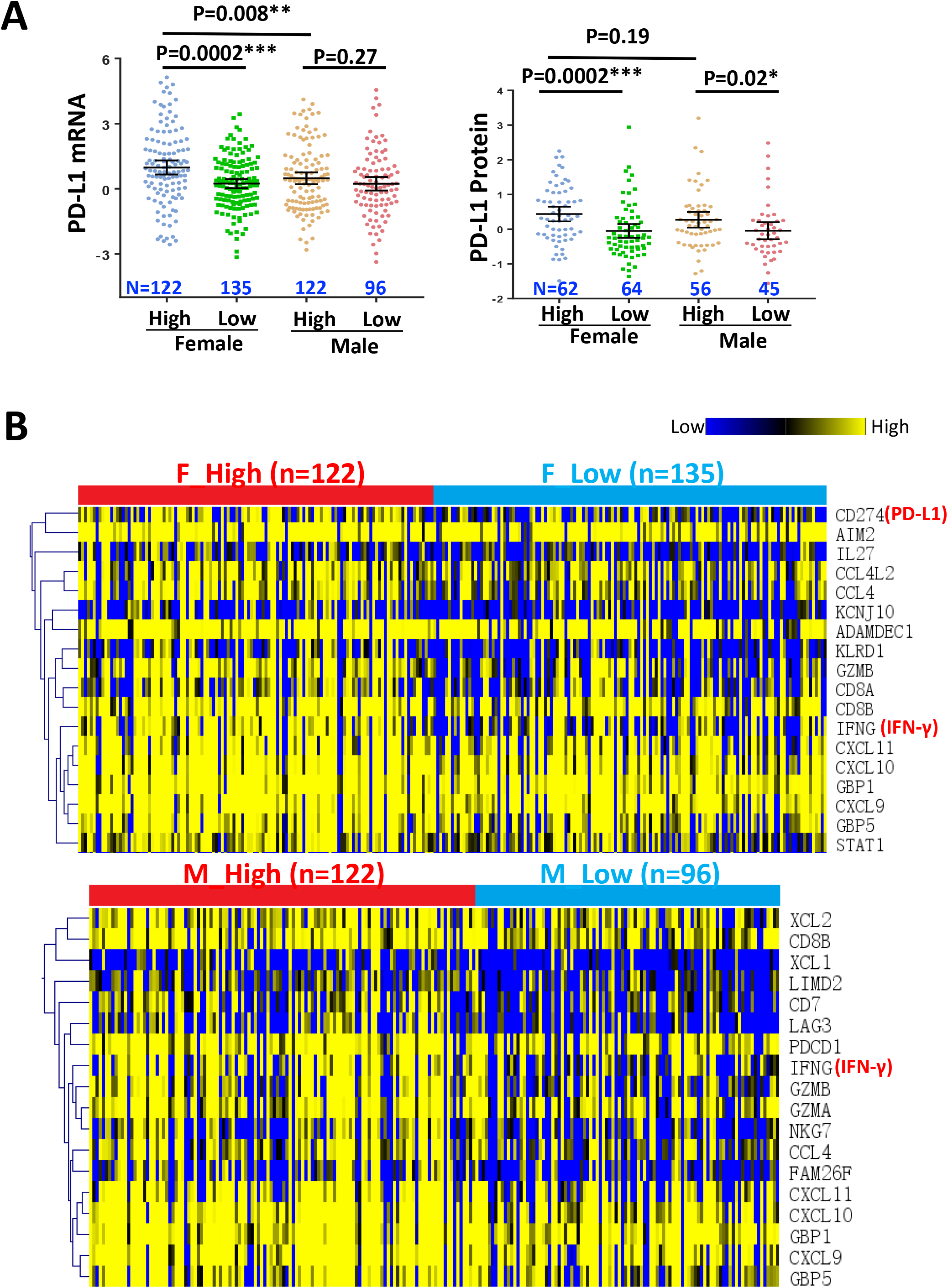
Gene signatures in *TCGA-LUAD*. **(A)** PD-L1 expression in the four patient groups stratified on Sex and TMB. **(B)** Hierarchical unsupervised clustering of genes differentially expressed in high TMB vs. low TMB groups in females (top) and males (bottom). t-distribution P value <0.005.

Finally, we examined tumor-infiltrating lymphocytes (TILs, including Tregs) and macrophages. Relative to the low TMB group, in females (but not males) with high TMB, TILs were enriched while the Treg subset of TILs and the M2 subtype of macrophages depleted (Fig. 6A, C, F). The Treg depletion could presumably contribute to the aforementioned T effector cell activation in these patients. The M2 macrophage depletion, perhaps resulting partly from KRAS mutations (35), might also enhance therapeutic effects. This is because macrophages, especially the M2 subtype, can strip anti-PD-1 antibody from T cells, and they also express PD-1 on the surface to sequester the anti-PD-1 antibody (37). In contrast to the Sex-specific effects described above, CD8 T cells and M1 macrophages were enriched in patients with high TMB irrespective of Sex (Fig. 6B, E). These data collectively indicate that for LUAD, a TME was robustly associated with TMB (and by inference, with therapeutic efficacy) only in females.

**Fig.6.**
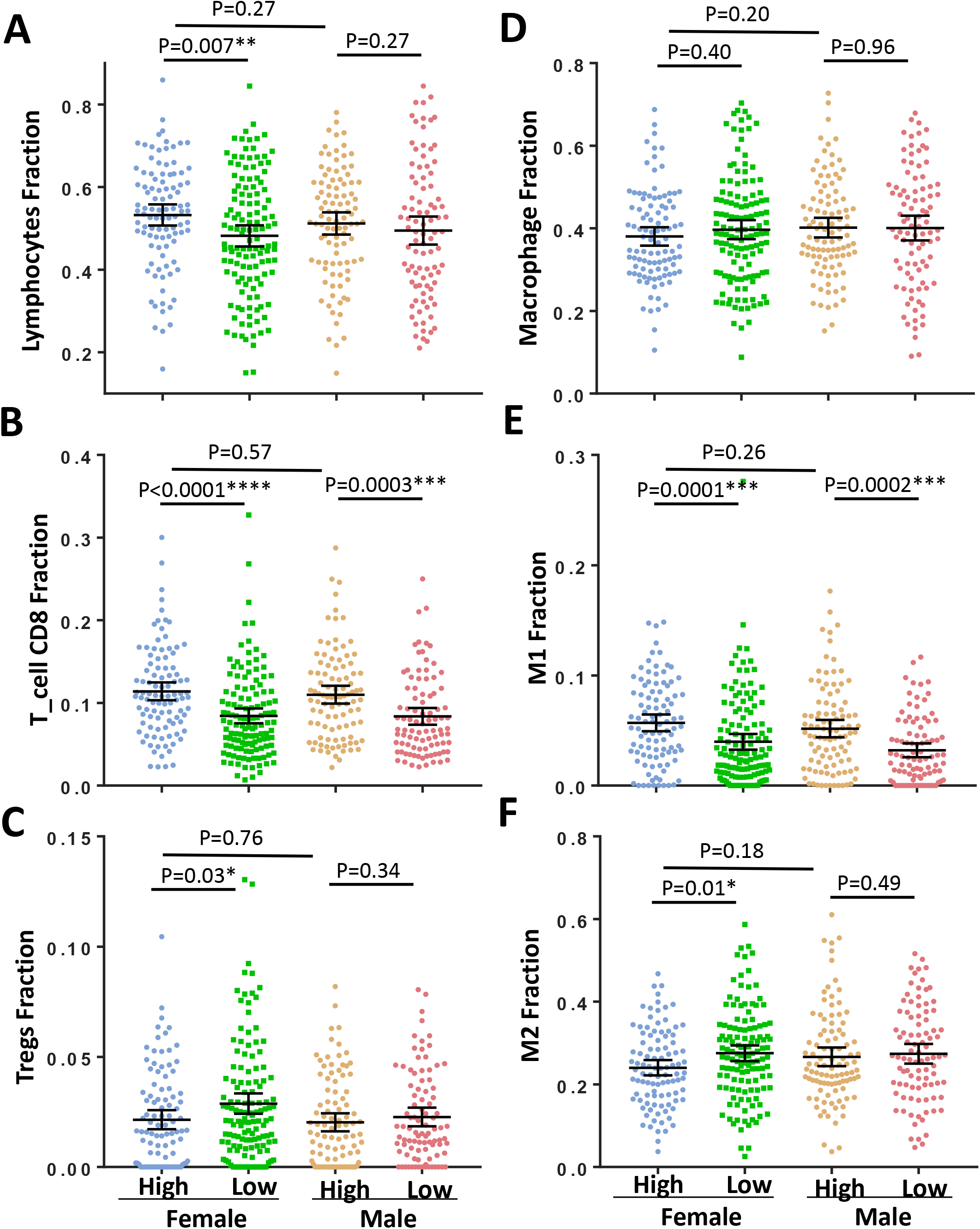
Tumor-infiltrating immune cells in the four patient groups in *TCGA-LUAD*. The cells analyzed were total lymphocytes (A) together with CD8 and Treg subsets (B-C), and total macrophages (D) together with the M1 and M2 subsets (E-F).

### Sex-dependent impact of biomarkers on PFS: multivariate regression analysis of the three immunotherapy datasets

Our analysis so far suggests that the 5 biomarkers examined (TMB, neoantigen burden, smoking signature, KRAS mutations and TME) can each predict therapeutic outcomes for LUAD in females. To corroborate and extend the conclusion, we found that these variables were highly correlated only in females, except that Tv/Ti was not significantly correlated with TMB/neoantigens in *Science-LUAD* perhaps due to its small sample size (Fig. S8A). Furthermore, in multivariate Cox regression analysis incorporating TMB, age, smoking status, and histology, TMB was independently associated with PFS in females (Fig. S8B). Of note, although males also had this trend, the trend was much weaker than females (Female: HR=0.11, P=0.007; Male: HR=0.06, P=0.06 in *Science-LUAD* cohort; Female: HR=0.20, P=0.002; Male: HR=0.90, P=0.8 in *Cancer Cell-NSCLC* cohort; Female: HR=0.28, P=0.002; Male: HR=0.88, P=0.7 in *JCO-LUAD* cohort; Fig. S8B).

## Discussion

Many biomarkers have been proposed to have predictive values for immunotherapy efficacy (5, 8). Here we report, for the first time, that at least for LUAD treated by anti-PD(L)1, five commonly used biomarkers (TMB and neoantigen burden, smoking signature, KRAS mutation, and TME) are robust only in females. Previous studies on the three immunotherapy cohorts all concluded that TMB/neoantigen had predictive values while discrepancies existed regarding Tv/Ti and KRAS (Table 1). Our data demonstrate that the predictive values of TMB and neoantigen burden reflected mainly the biomarker performance in females, and that the discrepancies regarding Tv/Ti and KRAS arose from confounding Sex effects, as the discrepancies vanished once the analysis was performed on patients stratified according to Sex (Table 1, cyanine shaded). As Sex is typically ignored in previous studies on immunotherapy, our work helps explain why patient responses to immunotherapy has been difficult to predict, at least for LUAD. It is clearly imperative to revisit immunotherapy datasets involving other forms of cancers/treatment regimen, in order to determine whether the conclusion on LUAD is generalizable.

One limitation of this study is that we could not analyze the predictive power of sex-based PD-L1 expression in three immunotherapeutic cohorts. In consideration of the poor quality of PD-L1 expression data, we abandoned it for further analysis. Although both PD-L1 expression and TMB are validated as biomarkers for immunotherapy response in phase III clinical trials (38, 39), TMB outperforms PD-L1 expression in NSCLC (1–3). Therefore, we mainly focused on sex-based TMB for immunotherapy in this study.

Multiple mechanisms might underlie the Sex-specific effects described above. For example, sex dimorphism has been observed in the human immune system (9). In particular, women are endowed with stronger innate and adaptive immune responses in adulthood, and are also more prone to autoimmune diseases (9), which might contribute to the favorable tumor immunotherapeutic microenvironment in the female LUAD patients with high TMB. Furthermore, females are more susceptible to tobacco carcinogens and hence more prone to KRAS mutations (11, 34), which might help explain the better outcome of PD-1 blockade, given that KRAS mutations might underlie PD-L1 upregulation and favorable tumor immunotherapeutic microenvironment (35). Of note, consistent with our finding, previous studies suggest KRAS mutations could increase the sensitivity to PD-1 blockade (1, 7), in contrast to other mutations such as EGFR mutation that has the opposite effect (4). Our study paves the way for future investigations into the mechanisms of sex dimorphism in the efficacy of LUAD immunotherapies.

Our study has implications for a recent report by Conforti and colleagues, based on the meta-analysis of 20 randomized controlled trials, that cancer immunotherapy is more effective on males, as measured by the survival hazard ratio (HR) (13). However, caveats in the meta-analysis have been pointed out, and contradictory results presented. In particular, it has been argued that HR (the relative hazard for immunotherapy versus control treatment) is not an adequate measure of therapeutic benefits and difficult to interpret clinically (40, 41); progression-free survival (PFS) should be compared, and for cohorts of melanoma patients on immunotherapy, 3-year PFS was not different between men and women (40). Neither could we detect Sex effect on PFS in the three NSCLC cohorts if we stratified the patients simply according to Sex (not shown; the three NSCLC cohorts were not included in the Conforti study). Note that in the Conforti study, the HRs in different NSCLC cohorts are quite variable. Specifically, five NSCLC cohorts treated with anti-PD-1 were included in the meta-analysis. For two of them, the HRs hardly differ between sexes (the differential being 0.04-0.05; Fig. 2), while the other three have much larger HRs (differential up to 0.42). The basis for this huge variability is unclear, partly because the datasets lack information about crucial biomarkers like TMB. Our data argue that if Sex does determine therapeutic efficacy in a particular type of tumor, it is only one of the determinants, and so other variables should be controlled for before the true role of Sex could be convincingly teased out.

## Acknowledgments

The authors thank the TCGA project and other groups for providing invaluable datasets for bioinformatic analysis. The authors also thank Lin-Li Yao for her important advice.

## Data Availability

All datasets generated for this study are included in the manuscript.

## Author contributions

MJ initiated and performed the analysis. TC supervised the project and wrote the manuscript with MJ.

---

**Fig. S1.**
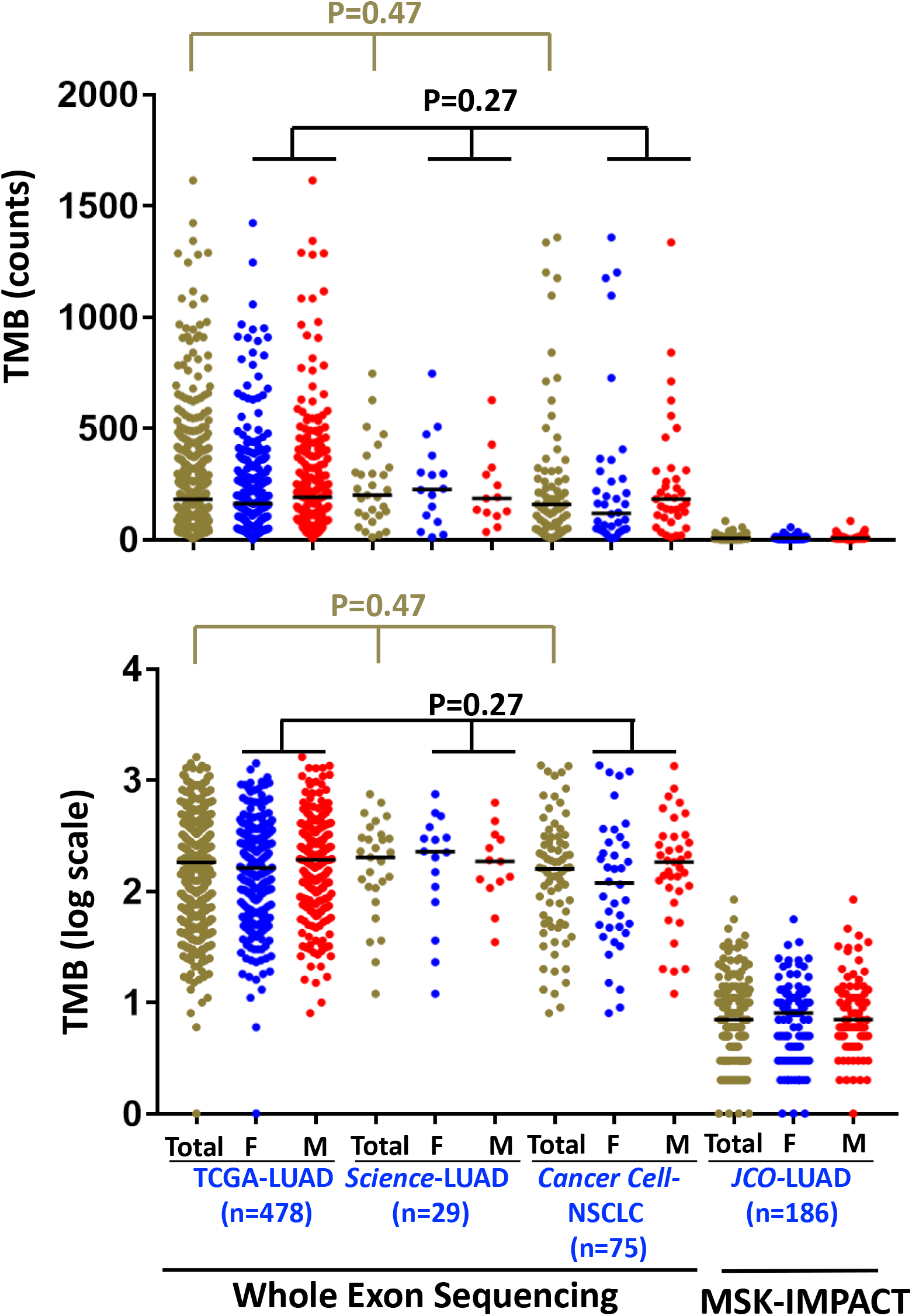
TMB in the 4 datasets used in the current study. Shown are the actual counts (top) and transformed values. Sex differences in TMB within dataset are not statistically significant.

**Fig. S2.**
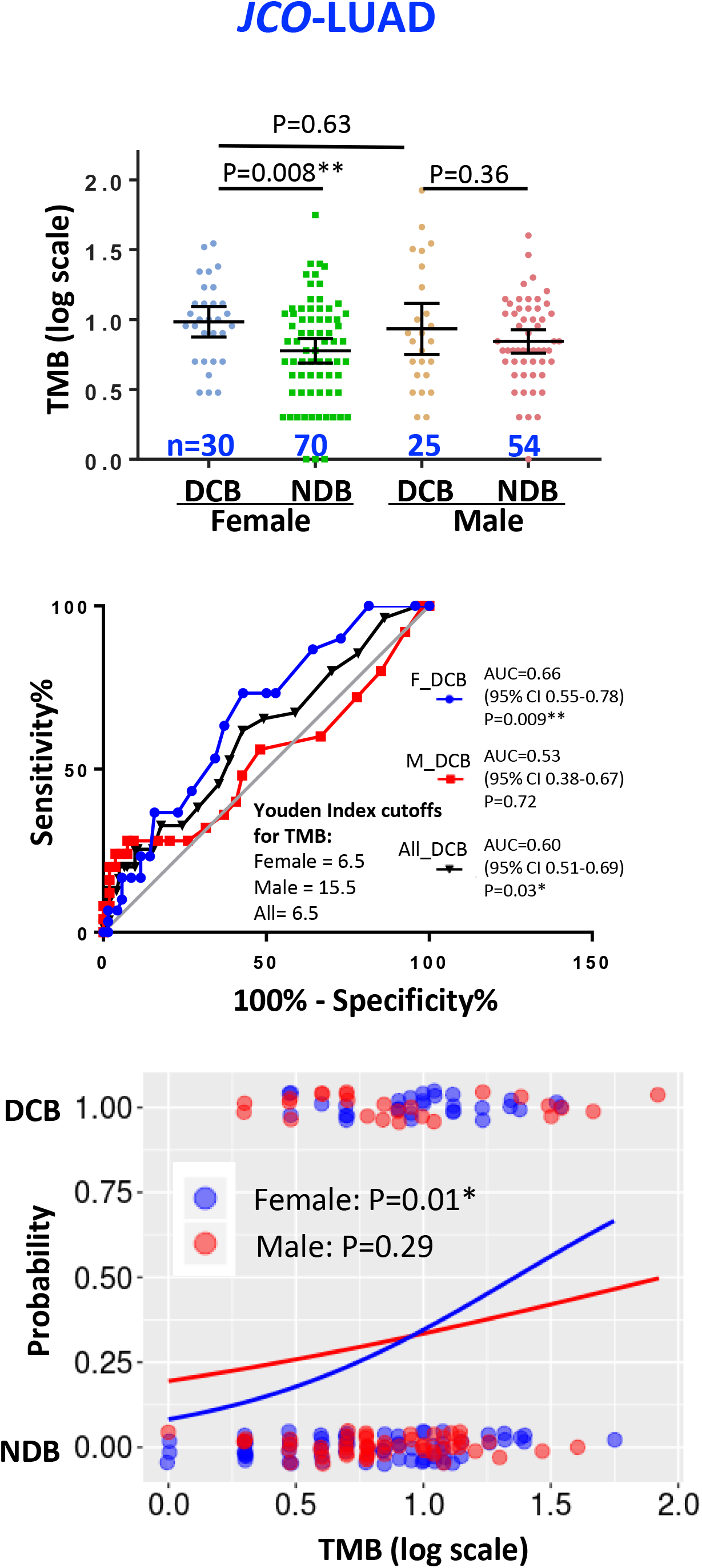
TMB correlation with patient responses in *JCO-LUAD*.

**Fig. S3.**
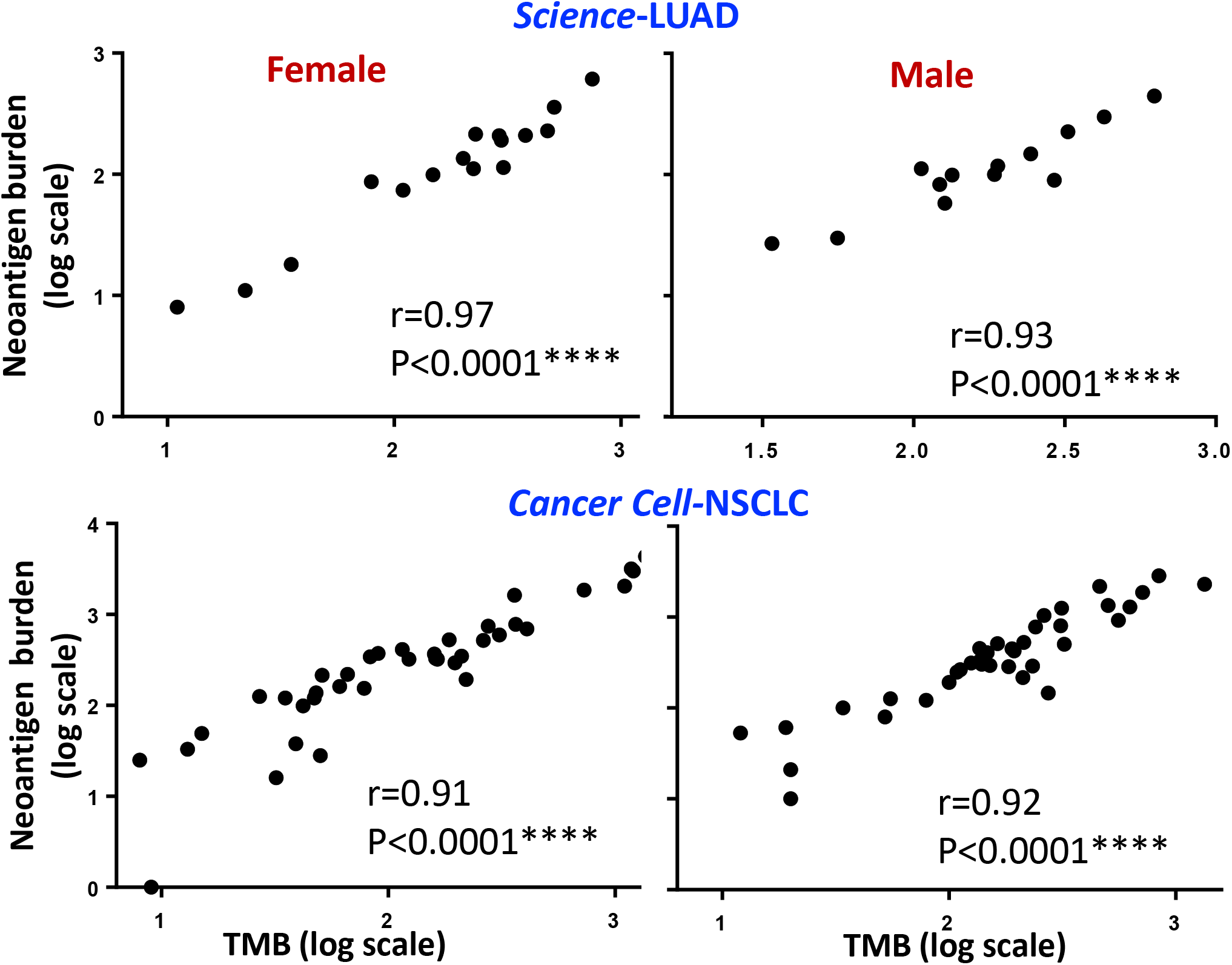
TMB correlation with neoantigen burden. r, Pearson correlation coefficient.

**Fig. S4.**
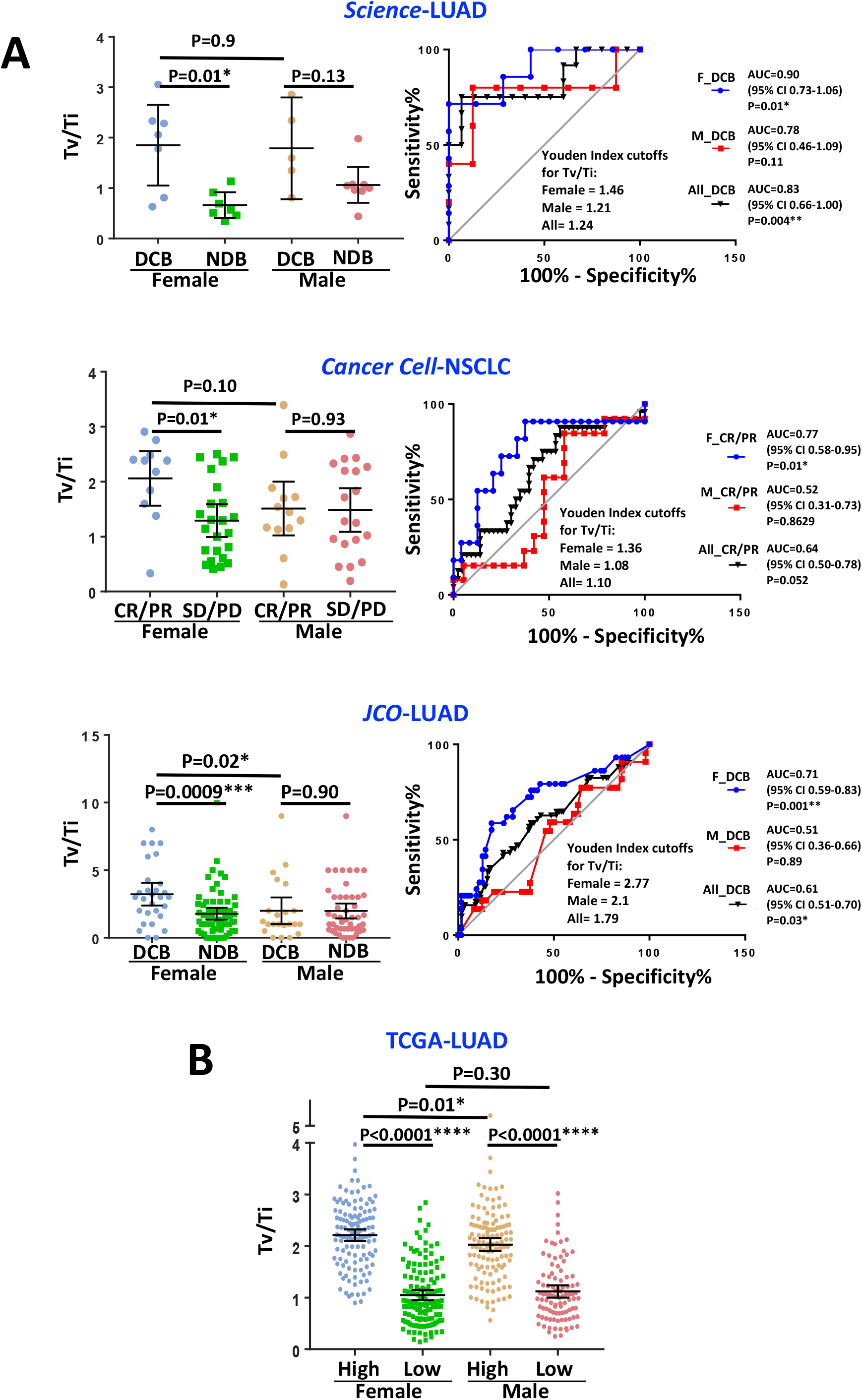
Sex effects on the predictive values of Tv/Ti in the immunotherapy cohorts. **(A) and *TCGA-LUAD* (B).** Patients in *TCGA-LUAD* were stratified on Sex and TMB.

**Fig.S5.**
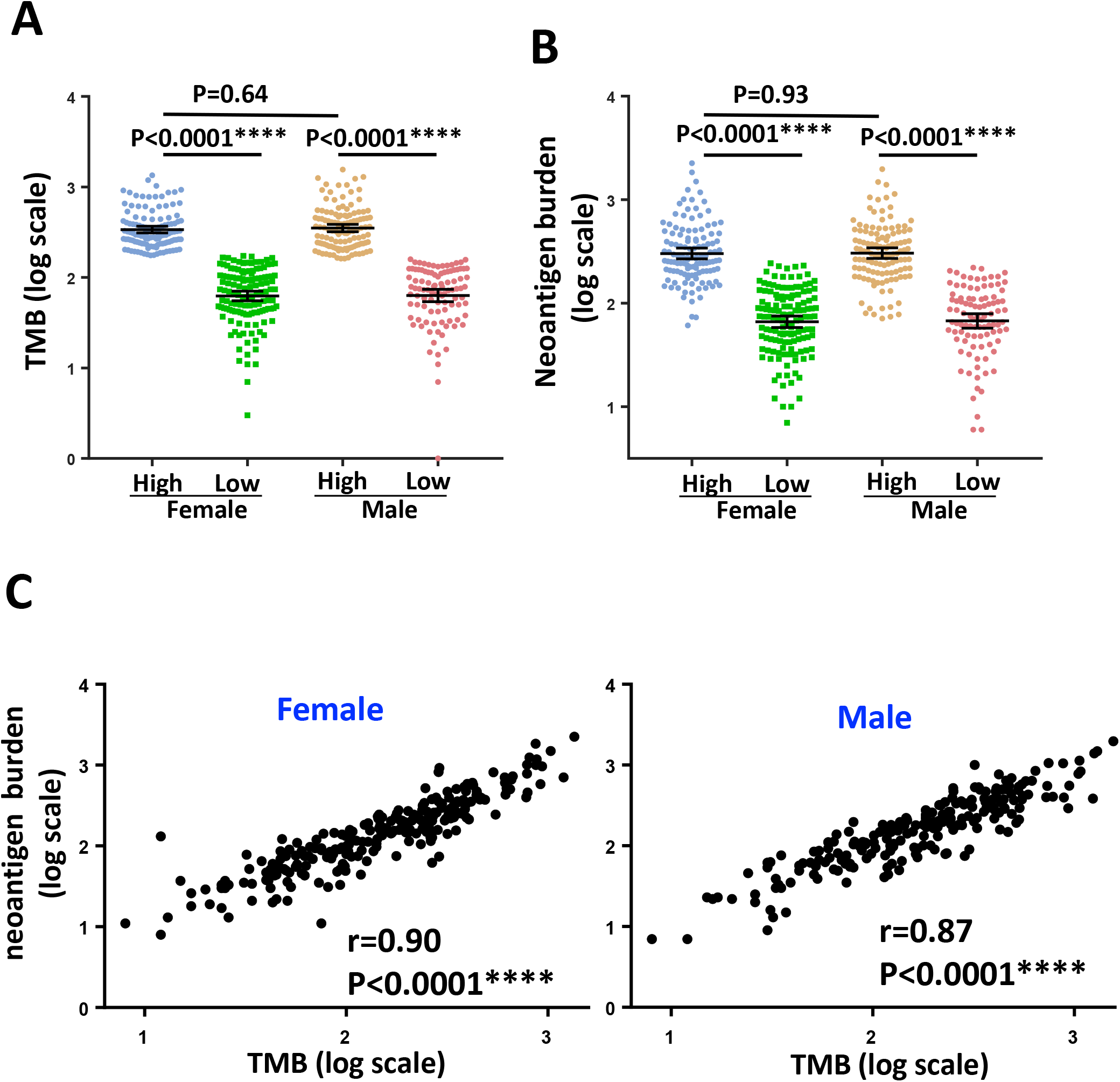
TMB and neoantigen burden in *TCGA-LUAD*. **(A)** Patient stratification into 4 groups based on Sex and TMB as in Fig. S4. **(B)** Neoantigen burden in the 4 patient groups. **(C)** Correlation between TMB and neoantigen burden. r, Pearson correlation coefficient.

**Fig. S6.**
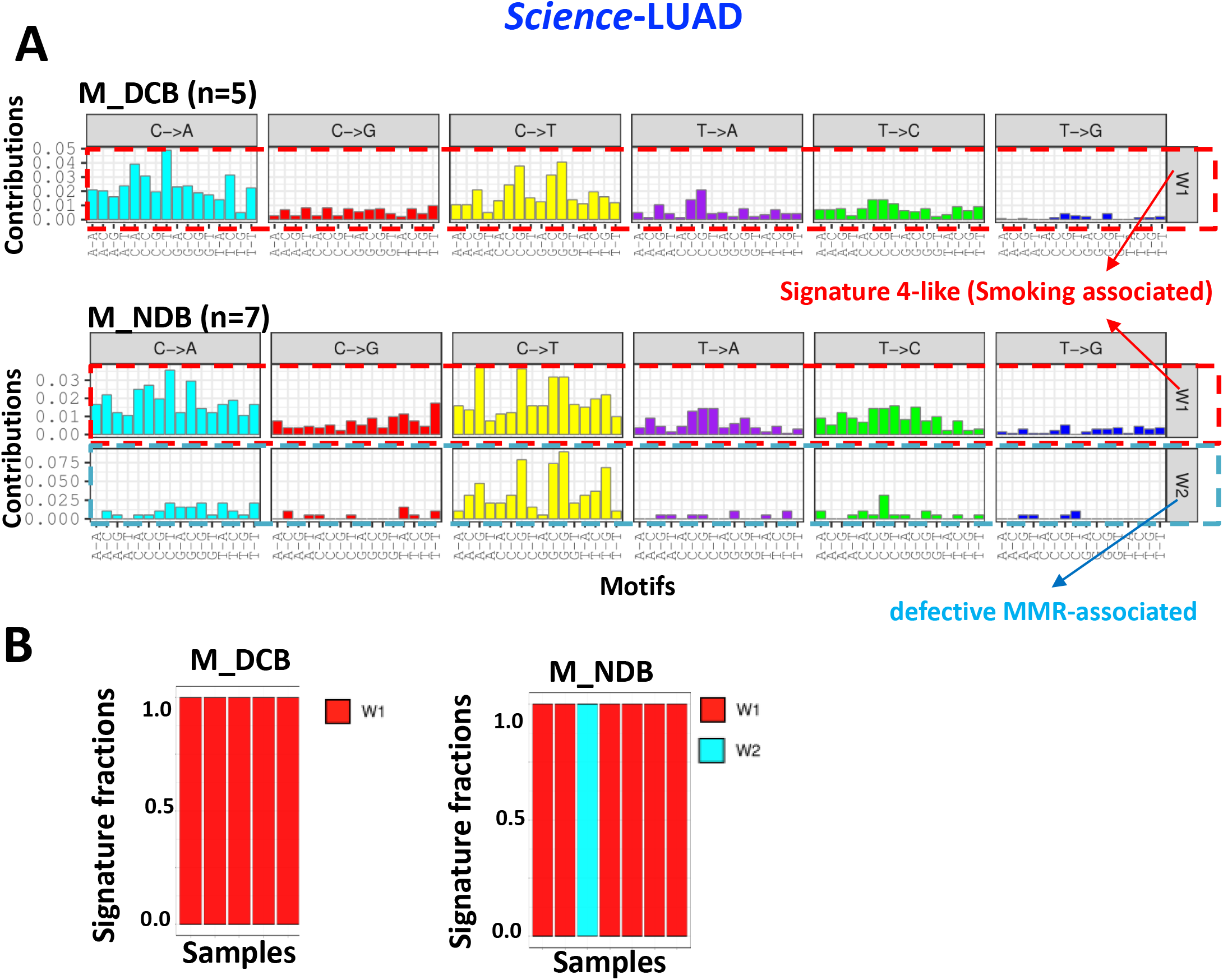
Mutational signatures in males in *Science-LUAD*. The smoking related signature was present in both DCB and NDB groups.

**Fig. S7.**
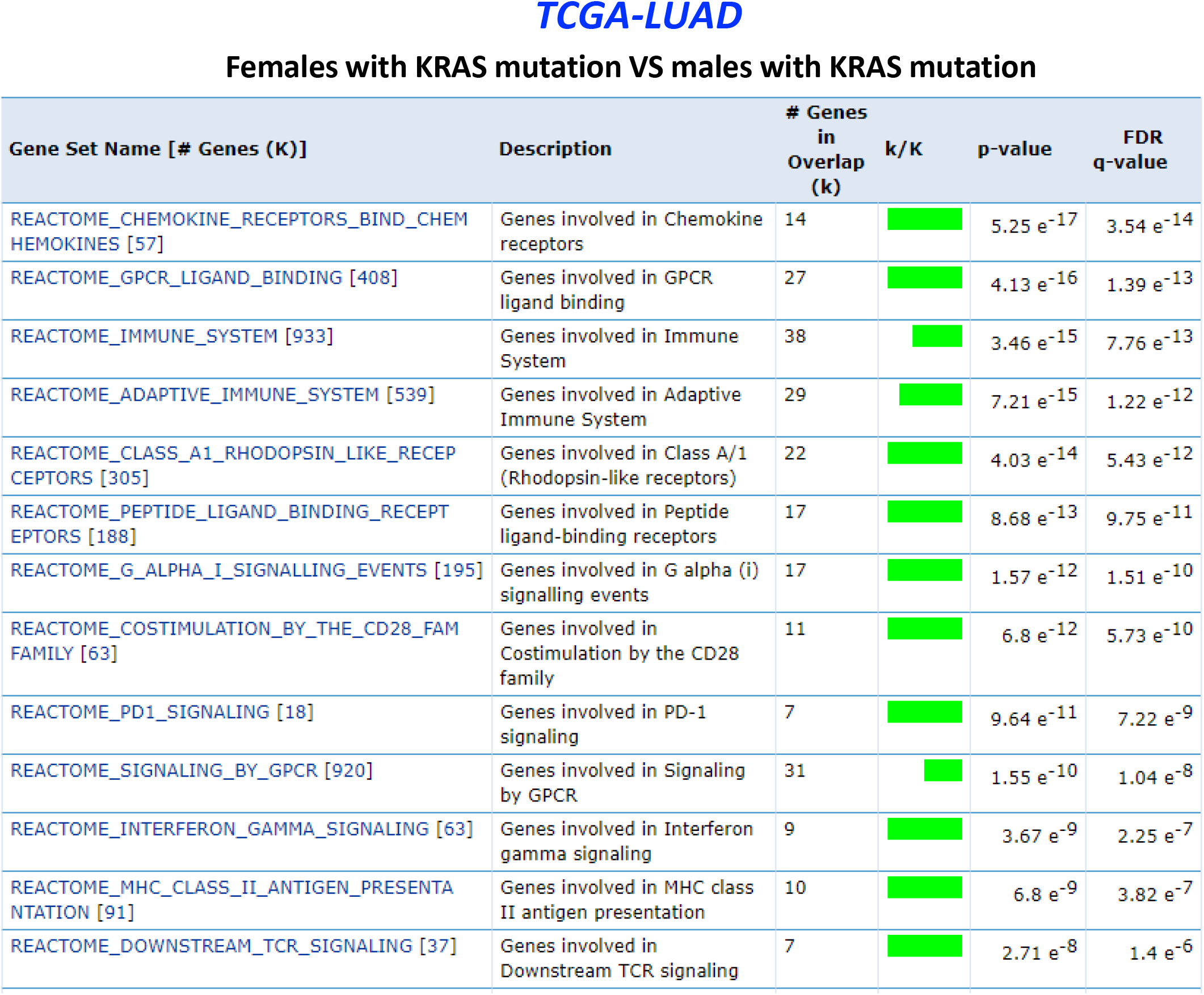
Gene set enrichment analysis revealing Sex-specific effects of KRAS mutations. Immune-related pathways were preferentially activated in females. Green bars indicate significant FDR q-values (< 0.05).

**Fig. S8.**
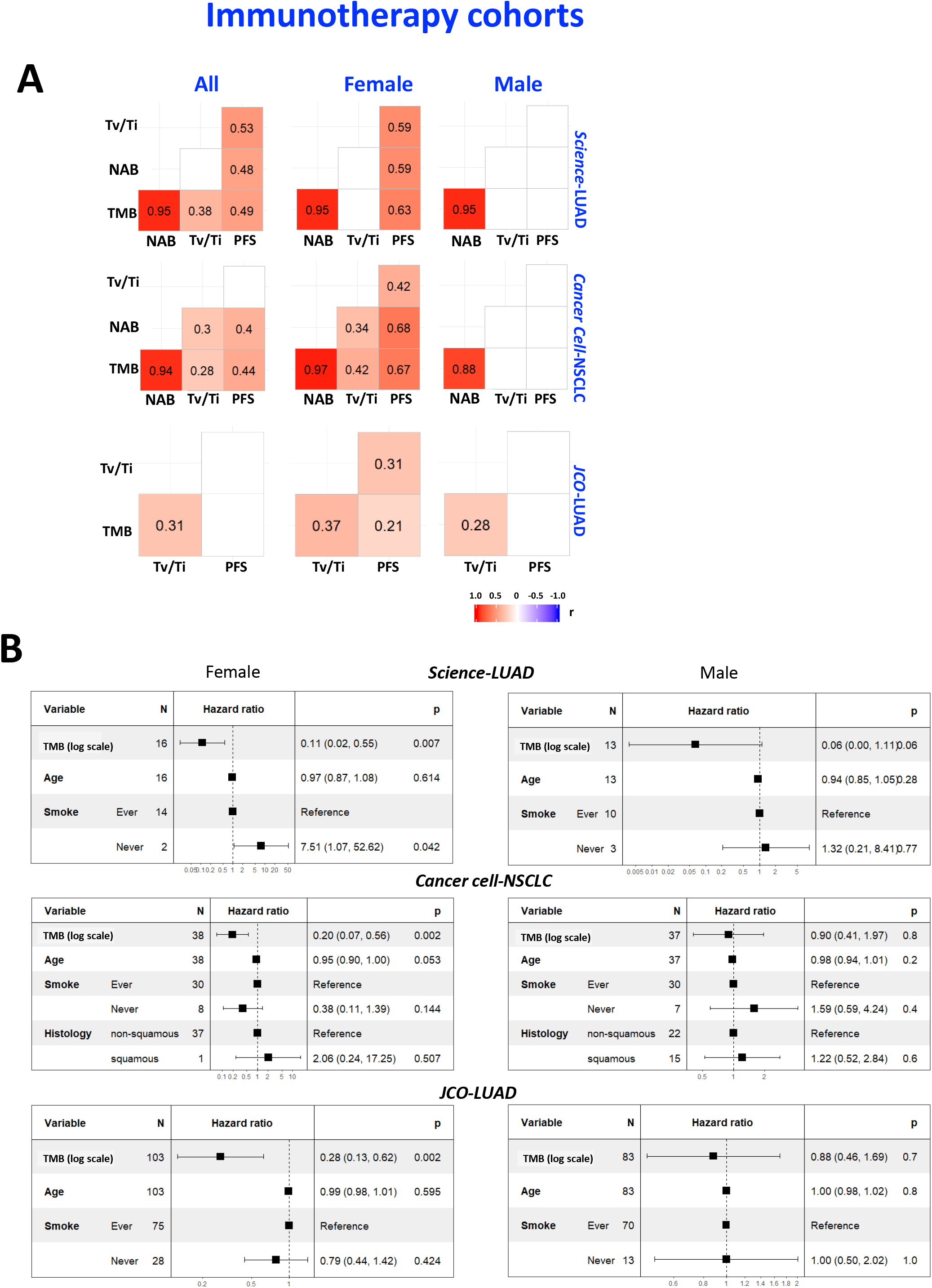
Multivariate Cox regression analysis. **(A)** Pairwise correlations between PFS and various biomarkers in males and females. Blank square indicated no significant correlation (P>0.05). The values are Pearson correlation coefficient (r). NAB, neoantigen burden **(B)** Multivariate Cox proportional hazards analysis of PFS in patients treated with immunotherapies demonstrating the hazard ratios (HR) for each covariates. Of note, neoantigen burden and Tv/Ti were not analyzed in Cox regression given their tight correlation with TMB.

